# Dissecting the membrane association mechanism of aerolysin pore at femtomolar concentrations using water as a probe

**DOI:** 10.1101/2024.01.18.576179

**Authors:** Tereza Roesel, Chan Cao, Juan F. Bada Juarez, Matteo Dal Peraro, Sylvie Roke

## Abstract

Aerolysin is a bacterial pore-forming toxin able to form transmembrane pores at the host plasma membrane of narrow internal diameter and great stability. These assets make it a highly promising nanopore for the detection of biopolymers such as nucleic acids and peptides. While much is known about aerolysin from a microbiological and structural side, its membrane association and pore-formation mechanism are not yet fully disclosed. Here, we studied the interaction of femtomolar concentrations of aerolysin and its mutants with liposomes in aqueous solution using angle-resolved second harmonic scattering (AR-SHS), in combination with single-channel current measurements. The measurements were so sensitive to detect electrostatic changes on the membrane-bound aerolysin induced by pH variation induced by the changes in the hydration shell of aerolysin. We reported for the first time the membrane binding affinity of aerolysin at different stages of the pore formation mechanism: while wt aerolysin has a binding affinity as high as 20 fM, the quasi-pore state and the prepore state show gradually decreasing membrane affinities, incomplete insertion and pore opening signature. Moreover, we quantitatively characterized the membrane affinity of mutants relevant for applications to nanopore sensing. This approach opens new possibilities to efficiently screen biological pores suitable for conducting molecular sensing and sequencing measurements, as well as to probe pore forming processes.

## INTRODUCTION

Pore-forming proteins are a class of proteins secreted by a variety of organisms usually involved in defense or attack mechanisms, targeting the plasma membrane^1^. The most extensively characterized pore-forming proteins are the bacterial pore-forming toxins (PFTs), which depending on the secondary structure defining the protein, have been classified as helical (α-) or beta-sheet (β)-PFTs^1,2^. Usually, pore-forming proteins are expressed as soluble proteins that subsequently oligomerize upon protease activation or membrane receptor binding, converting to a transmembrane pore.

Aerolysin, a β-PFT produced by *Aeromonas sp*., is the founding member of a large superfamily that spans all the kingdoms of life^1,3,4^. Aerolysin is expressed as an inactive precursor, proaerolysin, which contains 4 distinct domains: domain 1 (in gray, Figure 1A) is involved in binding N-linked oligosaccharides while domain 2 (in blue) is a glycosyl phosphatidylinositol (GPI)-anchored binding region, domain 3 (in yellow, called stem loop) is responsible for the oligomerization process and domain 4 (in green) contains a C-terminal peptide (CTP, in red) required for folding into the soluble monomeric form^5^. Proteolysis of the CTP allows aerolysin to oligomerize in a heptameric ring-like complex that inserts into the target membrane to form the pore via different intermediates (namely prepore, post-prepore, and quasi-pore intermediates, Figures 1B-D)^6^. These are defined by a different length and completion of the β-barrel which is formed after the stem loop undergoes a conformation change forming the β-barrel pore upon oligomerization. First, a transient prepore state is formed which is soluble and was captured *in vitro* only by the stabilizing Y221G mutation; this state is characterized by two concentric beta barrels, held together by hydrophobic interactions^4^ (Figure 1B). Next, in the series of events putatively leading to pore formation, the inner β-barrel fully extends towards the membrane passing from a quasi-pore state captured by the stabilizing mutation K246C-E258C (Figure 1C), and ending to the mature aerolysin pore (Figure 1D). All these different states have been characterized and their structure is known at high resolution by a combination of molecular modeling and cryo-EM experiments^4,7^, which allow to obtain a crude understanding of the sequential steps leading to pore formation.

**Figure 1:**
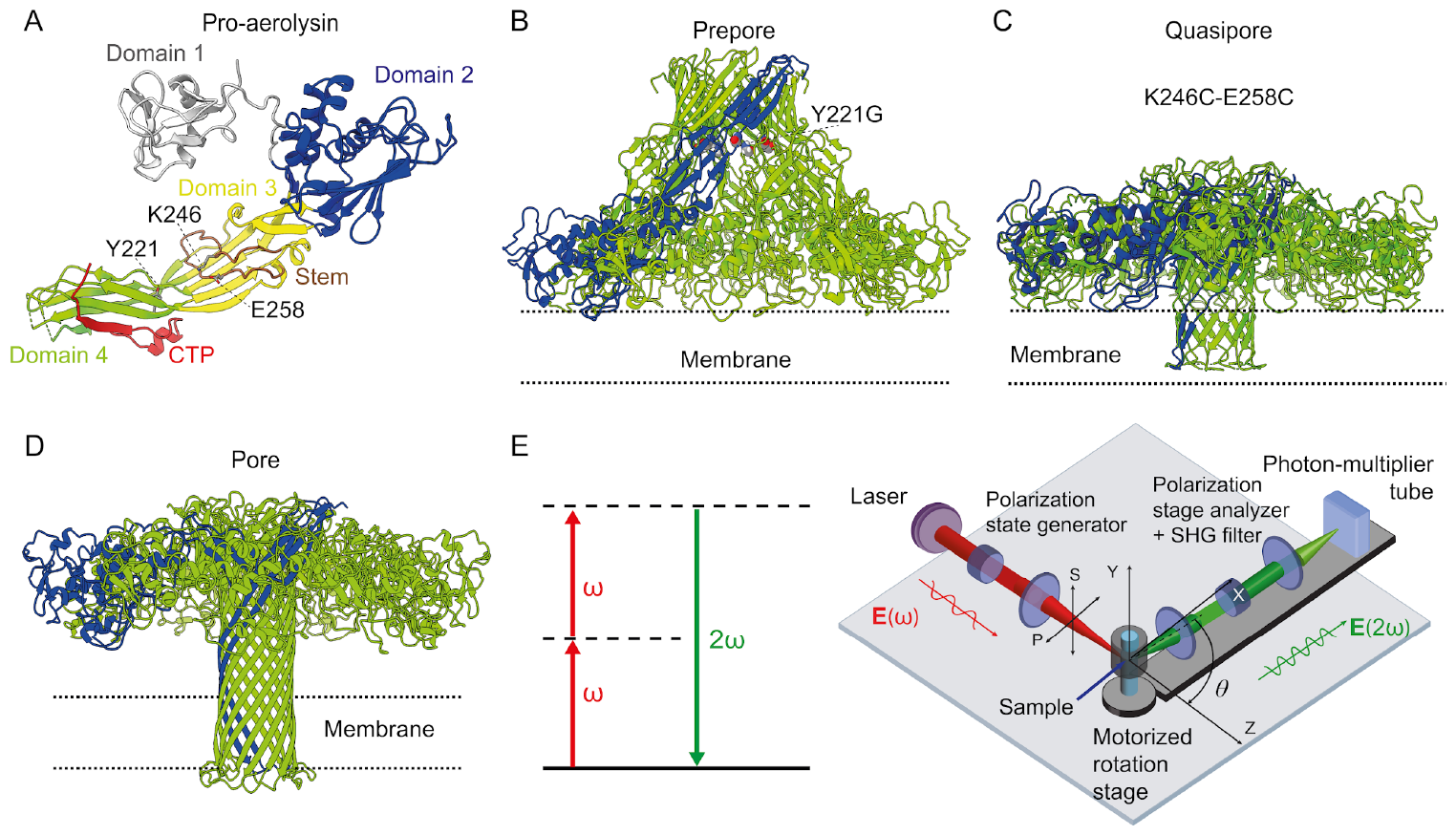
Structure of aerolysin in different states and the AR-SHS setup. **A**. Structure of monomeric pro-aerolysin (PDB:1PRE). Domain 1 in gray, Domain 2 in blue, Domain 3 in yellow and Domain 4 in green. Illustration of the aerolysin structure in pre-pore (**B**), quasi-pore (**C**), and pore (**D**) states, and labels of the various mutations with localization in panel A. **E**. Energy level scheme and a sketch of the AR-SHS experiment. P(S) refers to the polarization state of the beam parallel (perpendicular) to the scattering plane. All measurements were recorded with all beams polarized in the horizontal plane. For the single angle experiments, the scattering angle θ was set to 45° corresponding to the angle with maximum scattering intensity.

Recently, aerolysin has received much attention not only because of its biological role but also its potential use in biotechnological applications^8,9^. Due to its stability, the easy incorporation into lipid bilayers and a narrow internal diameter of the pore lumen (∼1 nm), aerolysin is nowadays one of the most promising biological nanopores, permitting the detection of several biopolymers such as nucleic acids, peptides, and oligosaccharides^10–14^. We and others have shown that point mutations within the pore can tune its electrostatics and steric properties for molecular sensing thereby achieving enhanced sensitivity and selectivity^15,16^. Meanwhile, we also observed that the mutations have an impact on pore formation efficiency. This is, in fact, crucial since without an efficient pore incorporation into the lipid bilayer and a strong lipid-pore interaction, nanopore experiments could not be conducted reliably. Therefore, to explore new biological or *de-novo* designed nanopore candidates, it would be highly desirable if one could estimate lipid-protein interactions at very low concentrations. Such a technique would be a useful tool to screen for pore mutants that bear stable and strong lipid interactions providing insights into the membrane association mechanism. It will be instrumental in identifying promising nanopore candidates to be used for molecular sensing experiments.

To better understand the complex aerolysin-membrane interplay we use a recently introduced method called high-throughput angle-resolved second harmonic scattering (AR-SHS)^17^. In this method, two femtosecond pulsed near infrared laser beams interact with an aqueous solution containing liposomes. A non-resonant second-order polarization is created in only those regions of the sample where there is an anisotropic distribution of anisotropic molecules. This polarization is essentially comprised of displaced charge that oscillates at twice the frequency of the incoming light, and it creates photons at this second harmonic frequency, which are detected. The AR-SHS experiment is illustrated in Figure 1E. It was recently shown that for liposome-water and other membrane systems, interfacial water molecules can be used as a contrast agent^18–22^. Thus, this technique is a label-free interface-specific method with molecular sensitivity towards the orientational distribution of interfacial water. Recent studies have shown that changes in the membrane water can help understand temperature induced phase transition^20^, and map different protein-membrane interactions such as the interaction of *α*-synuclein and the aqueous environment of liposomes^21^. It has also been shown that SHS enables the retrieval of the protein-membrane binding constant, which was exemplified by the interaction of perfringolysin O (PFO) with liposomes^23^. These measurements show the ability of SHS to ultra-sensitively detect protein-liposome interactions in an aqueous solution in the fM - pM range, corresponding to a single transmembrane protein being bound to a single liposome, using small volumes of protein solution.

Here, we used this approach to quantify the interaction between aerolysin pore-forming proteins and lipid membranes (Figure 1). Our results clearly show that SHS can capture pH-dependent changes in the surface charge of the aerolysin cap region. Combining the ultrasensitive SHS measurements with single-channel current recording experiments, we quantitatively analyzed the binding affinities of different aerolysin mutants to the lipid membrane. We estimated a binding affinity of 20 fM for wt aerolysin in a buffer solution with pH 7.4. Furthermore, we extracted the dissociation constants of two mutants – the first one mutated at the cap region (i.e., pore entry) - R220A and the second one mutated at the stem region (i.e., pore exit) - K238N. We observed a decrease of around two orders of magnitude between these two mutants, with K_d_ going from 10^−14^ M to 10^−12^ M for K238N and R220A, respectively. Additionally, we compared the binding behavior of mutation Y221G, which is known to block the transition from prepore to pore (Figure 1B), and of mutation K246C-E258C (Figure 1C), which forms a quasi-pore. No complete pore opening was observed for either of the mutants. However, the quasi-pore exhibited binding behavior similar to the wt aerolysin, but with a binding affinity more than an order of magnitude smaller.

## RESULTS AND DISCUSSION

### Surface charge detection upon pH changing conditions

Figure 2A shows the normalized SH intensity (*S*) at the angle with the maximum intensity (*θ*_*max*_ = 45°) as a function of the pH of the aqueous solution containing liposomes and 5 × 10^−10^ M of wt aerolysin. The normalized SH intensity is the SH intensity of the sample from which we subtract the SH intensity of the bulk solution and we normalize this value by the incoherent SH contribution of bulk water (more details are provided in Methods). The SHS patterns for pH 4 and pH 9 are shown in Supplementary Figure 1. In this type of experiment, SHS reports on the orientation of water in the interfacial region, whereby a high orientational directionality leads to a high SH response, and a vanishing orientational directionality (isotropy) leads to a vanishing SH response. Since charge-water dipole interactions are the main driver of orientational directionality and can be used to determine the electrostatic surface potential^18^, the SH response is very sensitive to the amino-acid charges of the protein (with the DOPC membrane being overall charge neutral). At pH 4, the normalized SH intensity, which relates the difference in water orientation to the response neat bulk water (Eq. 1, Materials and methods), is the lowest at 0.02. At pH 4.5 the normalized SH intensity sharply increases by 0.06, and keeps gently increasing until pH 8, after which it rapidly increases to 0.2. The electrostatic potential on the surface of the extracellular part of the wt aerolysin is reported in Figure 2B, as the environment changes from acidic to basic. This result highlights how the protonation state of the amino acids present in the cap domain significantly changes with pH, producing a mostly positively charged pore at pH 4 and a negatively charged one at pH 9. It is interesting to notice that significant variations of *S* happen in correspondence to the pKa values of the relevant titratable residues (i.e., Asp/Glu, His and Arg/Lys) within the pH range explored. Indeed, the cap domain consists of seven monomers, each of which contains 28 aspartic acids, 12 glutamic acids, 17 arginines, 6 histidines and 16 lysines. When the pH is lower than 4, Asp and Glu residues are mostly protonated, which reduces the negative charge of the pore cap domain (Figure 2B, pH 4). Going from pH 4 to 5, the net charge of the cap domain becomes more negative. At pH 6, histidines start becoming deprotonated which reduces the positive charge of the pore. At pH 9, arginines and lysines start to be deprotonated, further increasing the negative charge on the cap domain.

**Figure 2:**
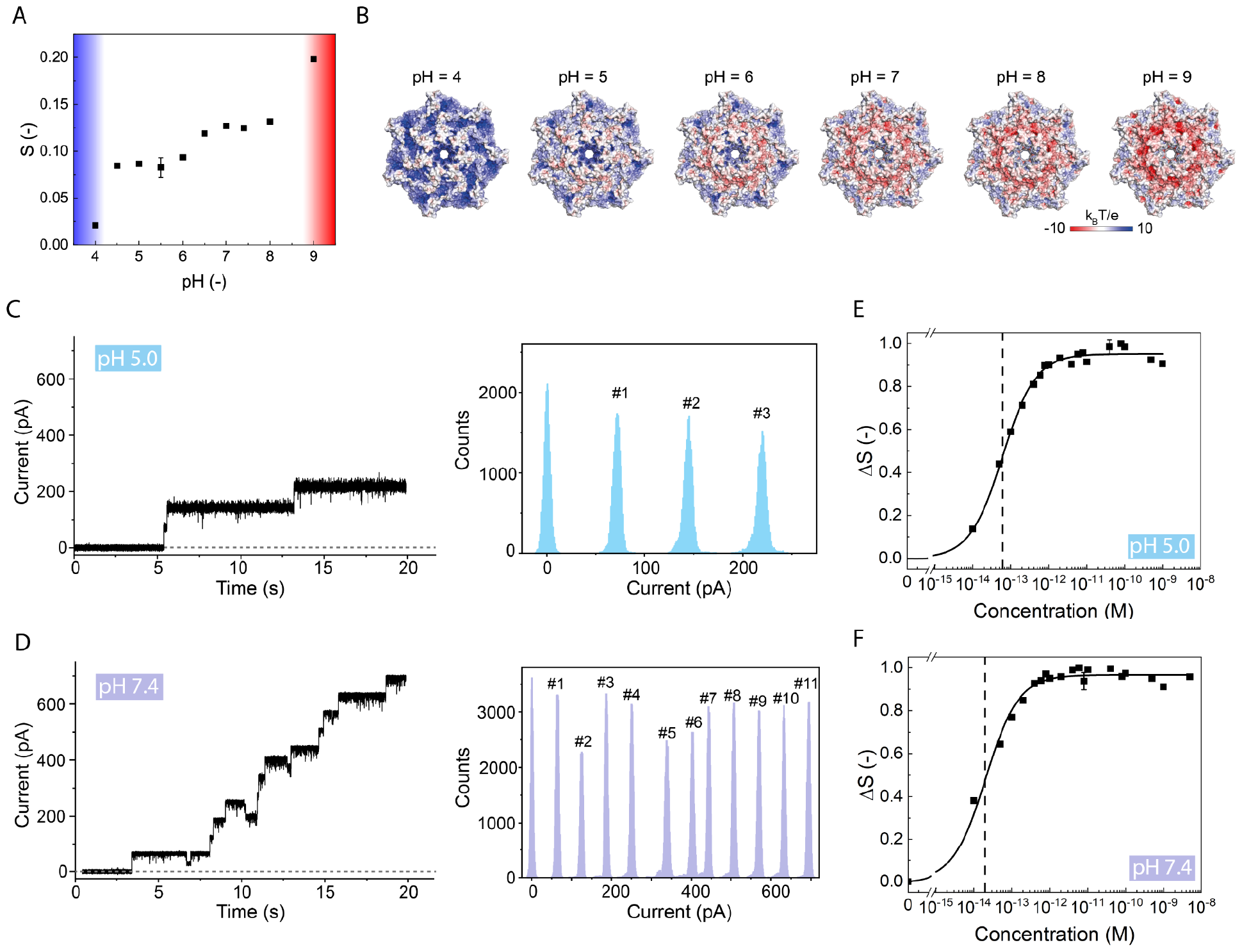
Surface charge changes and aerolysin-membrane binding affinities for different pH. **A:** Second harmonic intensity difference relative to bulk water (S) at the scattering angle with maximum intensity (*θ*_*max*_ = 45°) vs pH of the aqueous solution. The measurements were performed with DOPC liposomes and wt aerolysin. The blue and red highlighted areas indicate the positive and negative surface charge on the cap region of aerolysin. **B:** Electrostatic potential mapped on the cross-sections of the aerolysin cap domain. **C, D:** A representative single-channel current recording traces of the aerolysin-DOPC free-standing membrane system in aqueous solution at pH 5 (**C**) and 7.4 (**D**), with the corresponding histogram current distribution showing a single pore incorporation. **E, F:** Normalized SH intensity difference (*Δ S*) at the angle with the maximum intensity (*θ*_*max*_ = 45°) vs wt aerolysin concentration in logarithmic scale at pH 5 (**E**) and 7.4 (**F**). The data are fitted using equation (2) giving a dissociation constant of *K*_*d*_ = (6.2 ± 0.4) × 10^−14^ M for pH 5 and *K*_*d*_ = (2.0 ± 0.2) 10^−14^ M for pH 7.4 (represented by dashed line). The error bars were determined as the standard deviation from 100 measurements for all the SHS measurements. SHS measurements were performed with a DOPC doped with 1% DOPS liposomes and with wt aerolysin.

Examining the trend in Figure 2A, we can experimentally see the same surface charge behavior through the orientation of the water molecules: As the liposomes for this experiment consisted of pure zwitterionic DOPC lipids, the charge of the membrane stayed neutral within the measured pH range. It was previously shown that DOPC liposomes in an aqueous solution have water molecules preferentially oriented with their hydrogens towards the membrane surface^18^. At pH 4, the inserted aerolysin is more positively charged, which forces some water molecules to orient their oxygen atoms towards the aerolysin surface, balancing out the effect of the water orientation induced by the DOPC membrane. This reorientation of water molecules causes the SH intensity to decrease. Between pH 4.5 and 8, the aerolysin cap domain is mostly neutral or slightly negative. Herein, the water molecule orientation is primarily due to the interaction with the DOPC membrane, resulting in hydrogens pointing towards the surface. Above pH 8, the aerolysin’s surface charge becomes strongly negative, and more water molecules reorient with their hydrogens towards the surface causing an increase in the SH intensity. Therefore, the SHS response is a very sensitive marker of the state of protonation of the amino acids of aerolysin. We expect that this to be a general trend for all transmembrane proteins possessing titratable moieties under variable pH conditions.

### Membrane binding affinity of wt aerolysin pores

The aerolysin-membrane binding was probed using SHS and single-channel current measurements at pH 5.0, and 7.4. In single-channel current recording experiments (Figures 2C and D), the membrane incorporation of an aerolysin pore into a DOPC free-standing membrane is followed by a rapid increase in current when a voltage is applied, which is proportional to the number of pores present in the membrane. For the aerolysin-DOPC membrane system in an aqueous solution at pH 5.0, three aerolysin pores are observed in a 20 s recording time, whereas for pH 7.4, many more pores are incorporated in the bilayer under the same conditions and equal recording time. Figures 2E and 2F show the coherent SH intensity difference *ΔS* as a function of the wt aerolysin concentration in the solution. *ΔS* is defined (Eq. 2, materials and methods) as an absolute value of the difference between normalized SH intensity of liposomes with the given concentration of aerolysin in the solution and liposomes without added aerolysin. It shows a rapid increase between 10 and 100 fM of aerolysin. This increase happens at lower concentrations for the system at pH 7.4 compared to the one at pH 5.0. At around 1 pM, the SH intensity starts to level off representing the saturation of the interactions, where no additional proteins can insert to the membrane. Using equation (2) we can obtain a wt aerolysin dissociation constant of *K*_*d*_ = (6.2 ± 0.4) × 10^−14^ M and *K*_*d*_ = (2.0 ± 0.2) × 10^−14^ M for pH 5 and pH 7.4, respectively. The single-channel current and SHS measurements together demonstrate that wt aerolysin has a slightly lower membrane affinity at acidic conditions compared to neutral pH, which is reflected by less efficient incorporation into lipid bilayers.

### Membrane binding affinities of aerolysin mutants

Being able to detect protonation changes of the amino acids of wt aerolysin, we aim to detect the changes caused by mutating amino acids in particular at the cap or the stem region. We use two single point mutations defined by distinctive steric hindrance and electrostatics^8^. It was shown that by replacing R220 at the cap region with alanine (R220A) the pore diameter becomes two times wider while the insertion efficiency is lowered^8^. A mutation at the constriction point in the stem region, where K238 is replaced with an asparagine (K238N) was shown instead to prolong the dwell time for DNA sensing with respect to the wt pore.

We investigated the insertion of these mutants in comparison with the wt aerolysin at pH 7.4, employing single-channel current recording and SHS measurements (Figure 3). From the single-channel current recordings at pH 7.4, we observe that two R220A pores and five K238N pores are formed on the membrane (Figures 3A and B, again in the same time window of 20 s). Using the same pores, we performed AR-SHS measurements to obtain the dissociation constant and compared it to wt aerolysin. We observe that in the case of R220A, the increase is more gradual than wt starting at less than 1 pM till 100 pM of the mutant concentration, where it levels off (Figure 3C). In the case of K238N, the SH intensity difference increases in a similar concentration range (10 fM-1 pM), but more gradually than in the case of wt. It saturates at around 1 pM, without a further change in SH intensity (Figure 3D). Using equation (2) we obtain dissociation constants of *K*_*d*_ = (2.3 ± 0.3) × 10^−12^ M and *K*_*d*_ = (4.6 ± 0.6) × 10^−14^ M for R220A and K238N, respectively. The extracted dissociation constant of the K238N at pH 7.4 is slightly higher than the dissociation constant of the wt aerolysin, revealing that K238N has a slightly less favorable interaction with the membrane, confirmed by less efficient incorporation into the membrane. The R220A dissociation constant is two orders of magnitude higher than wt aerolysin, reflecting the fact that it is less effective at creating pores in the membrane as also confirmed by the current measurements (Figure 3A). Comparing the number of aerolysin pore incorporations in the free-standing bilayer with the dissociation constants obtained from SHS shows that the two techniques combined return a consistent and quantitative description of the membrane association process.

**Figure 3:**
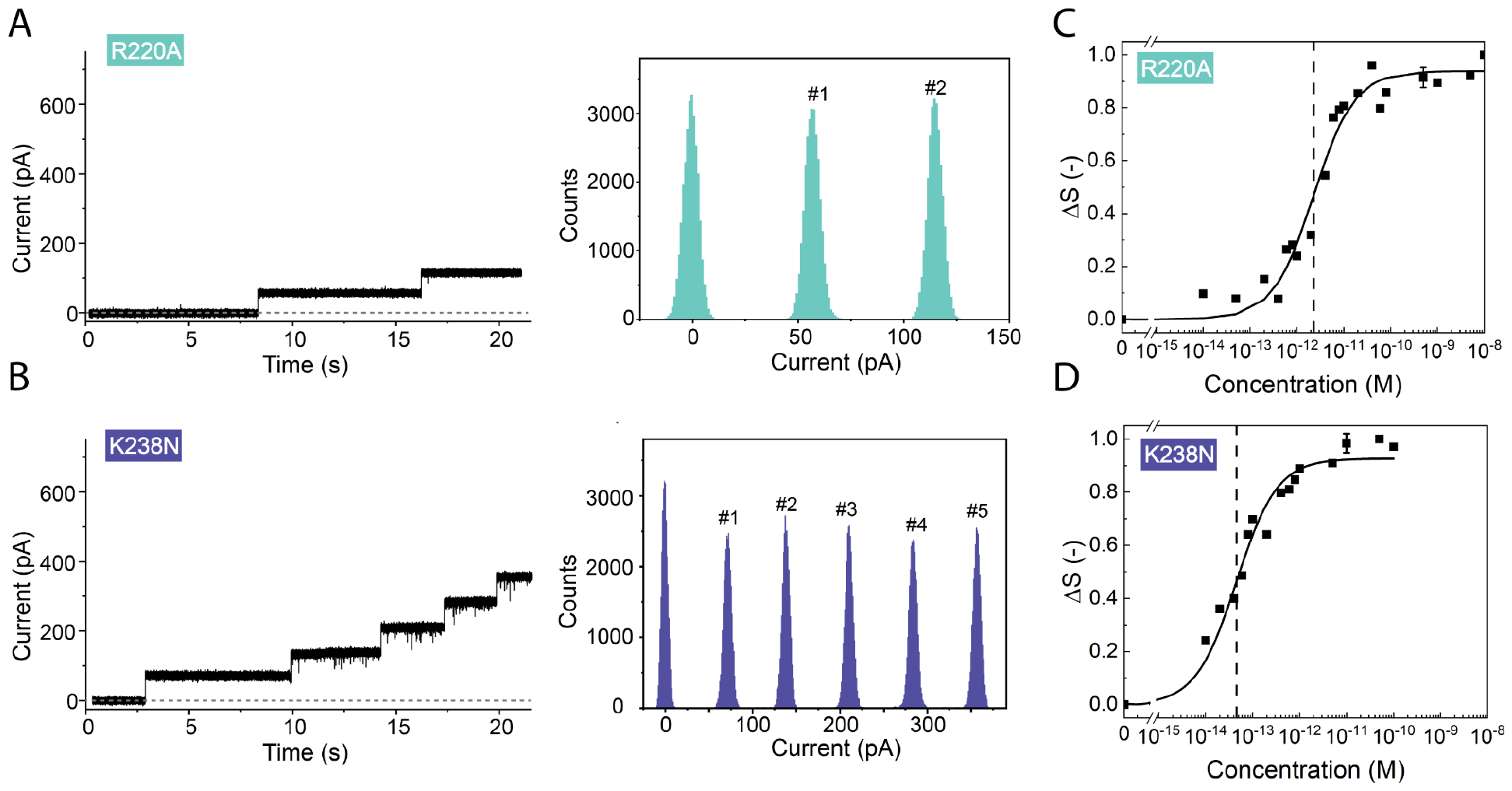
Binding of aerolysin single-point mutants to a lipid membrane. **A, B:** A representative single-channel current recording measurement of R220A (**A**) and K238N (**B**) measured with a DOPC free-standing membrane at pH 7.4. **C, D**: SH intensity difference (*Δ S*) at the angle with maximum intensity (*θ*_*max*_ = 45°) vs R220A (**C**), and K238N (**D**) concentration in logarithmic scale at pH 7.4. The data are fitted using equation (2) giving the dissociation constant of *K*_*d*_ = (2.3 ± 0.3) 10^−12^ M and *K*_*d*_ = (4.6 ± 0.6) 10^−14^ M for R220A and K238N, respectively (the dashed line represents the dissociation constant). These data were measured with DOPC doped with 1% DOPS liposomes. The error bars were determined as the standard deviation from 100 measurements for all the SHS measurements.

### Dissecting membrane association throughout the pore-forming process

To dissect more directly the mechanism of pore formation we observe how different pore intermediates (trapped by selected mutations Y221G and K246C-E258C, Figures 4A and B) interact with the membrane. Y221G pores are known to form a prepore state preventing the pre-stem loop from moving away from the five-stranded β-sheet in the same domain and therefore block the hemolytic activity of the toxin^7,24^. However, could be able to transiently interact with the target membrane although it is fully hydrophilic^24^. The K246-E258C mutations block instead the pore formation in a later stage of the formation of full transmembrane β-barrel due to the formation of a disulfide bridge between the mutated residues 246 and 258^5^.

**Figure 4:**
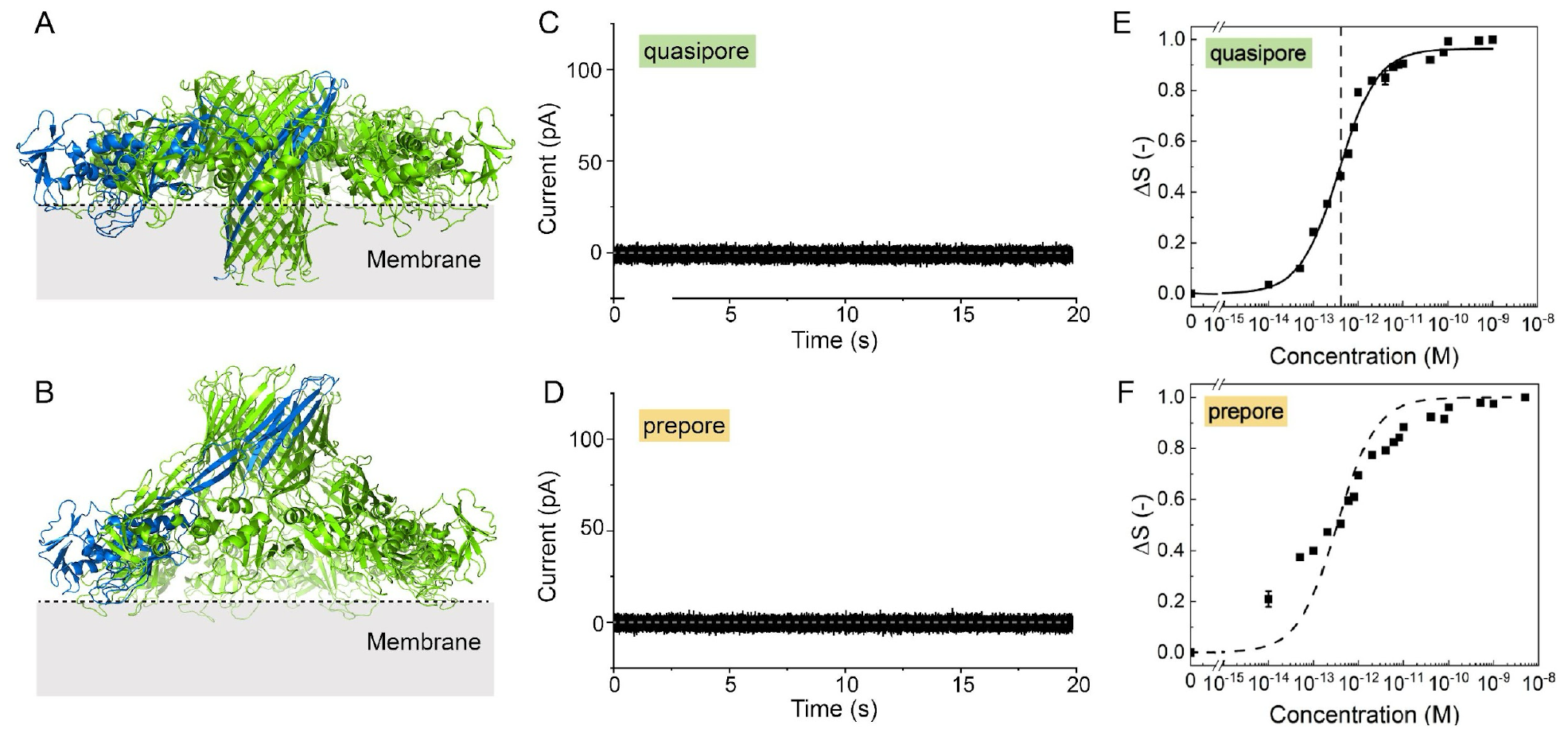
Membrane binding and conductivity of aerolysin mutants blocked at different stages of pore formation. **A, B:** Structural model of the aerolysin quasipore (**A**) and prepore (**B**). **C, D**: A representative single-channel current recording measurements of quasipore (**C**) and prepore (**D**) at pH7.4 on DOPC free-standing membrane. **E, F:** SH intensity difference (*Δ S*) at the angle with the maximum intensity (*θ*_*max*_ = 45°) vs K246C-E258C (**E**), and Y221G (**F**) concentration in logarithmic scale. The data are fitted using equation (2) giving a dissociation constant of *K*_*d*_ = (4.1 ± 0.3) × 10^−13^ M for the K246C-E258C (quasipore). These data were measured with DOPC doped with 1% DOPS liposomes at pH 7.4. The error bars were determined as a standard deviation from 100 measurements for all the SHS measurements.

Figures 4C and D both show a zero value for the whole time interval of the single-channel current measurement proving that there are no (or very few) pores inserted in the membrane. The SHS measurements provide insight into the aerolysin-membrane interaction. The SH intensity difference (*Δ S*) as a function of the concentration of the aerolysin mutants at pH 7.4 is depicted in Figures 4E and F. For the quasipore, *ΔS* is increasing from 10 fM to 10 pM where it saturates. The extracted dissociation constant for the quasi-pore is *K*_*d*_ = (4.1 ± 0.3) × 10^−13^ M, thus more than one order of magnitude higher than that of wt aerolysin (Figure 4E). The molecular interpretation of these results is that quasi-pores can bind efficiently to the membrane as they have an almost complete β-barrel that can insert into the membrane. As this is shorter compared to mature aerolysin pores the binding affinity is decreasing with respect to wt, moreover the barrel is not able to fully pierce the membrane preventing the formation of conducting pores. The combination of these results implies that the K246C-E258C mutant is inserted in the membrane, but not able to transition into the mature pore state, as shown previously^24^. In the case of Y221G, *ΔS* is increasing, but the increase does not follow the same trend as for the wt or previously mentioned mutants (Figure 4F). The fitting procedure fails indeed to work on this dataset, as indicated by the dashed line in Figure 4F. This could mean that the mutant is interacting with the membrane, but is not fully anchored to that, having only a transient membrane association behavior. The absence of detectable current on the single-channel recordings agrees with this assessment: No conducting pores are observed for the prepore mutant-membrane interaction. This is consistent with the fact that Y221G prepore mutation is known to form fully soluble pores and the barrel region is only partially folded, its hydrophobic stem region being still protected (Figure 4B).

Thus, by combining two complementary methods, one quantitative method reporting on water orientation (due to protein charging) that can be translated into protein-membrane interactions, and the other able to characterize single pore conductivity, we characterized the membrane association behavior of aerolysin pores. This approach can be generalized for other transmembrane proteins.

## CONCLUSIONS

In summary, using single-channel current experiments on free-standing membranes and AR-SHS measurements on aerolysin-LUV interactions in aqueous solutions, we showed that different mutants of aerolysin have different types of interactions with the membrane, leading to various degrees of pore formation and targeted insertion. Thanks to the high interface sensitivity of SHS, we were able to observe a change in the surface charge on the cap region of aerolysin bound to the LUV at different pH values of the aqueous solution. The charge of the cap region changes from positive at pH 4 to highly negative at pH 9 consistently with the predicted electrostatic properties of the pore. This provides a unique method to assess the electrostatic properties of surface of engineered biological nanopore applications which is very important for nanopore applications. Very recently, it has been demonstrated that the engineering of nanopore surfaces can induce a strong electroosmotic flow that has the advantage to capture all kinds of biomolecules regardless of their charges^25^.

By combining SHS and single-channel current measurements we compared the binding affinities of wt aerolysin at different pH. We extracted the dissociation constant of *K*_d_= (6.2 ± 0.4) × 10^−14^ M and *K*_*d*_ = (2.0 ± 0.2) × 10^−14^ M for pH 5.0 and pH 7.4, respectively. This is two orders of magnitude higher than the interaction of PFO with cholesterol-rich lipid membrane^23^. Additionally, we studied interactions of aerolysin mutants, R220A and K238N, and could recapitulate their distinct properties for nanopore sensing. We observed a difference of around two orders of magnitude between those two mutants, going from 10^−14^ M to 10^−12^ M for K238N and R220A, respectively. Lastly, we examined the different binding behavior of aerolysin mutants blocked at different stages of transition from prepore to pore. The quasipore was binding to the liposomes with a dissociation constant of *K*_*d*_ = (4.1 ± 0.3) × 10^−13^ M. No open-pore was observed for either quasipore or prepore mutants in single-channel current measurements, demonstrating the extreme sensitivity of the SHS technique which could detect as low as 10 fM of aerolysin in the aqueous solution with liposomes.

Having the possibility of using low volumes (as low as 10 µL of the final sample), low protein concentrations, and with the possibility to vary the temperature, pH, or membrane composition, the SHS method is able to distinguish which aerolysin pores are efficiently incorporated into the lipid bilayer providing the opportunity to dissect the molecular features of membrane association along the pore formation pathway. As shown for aerolysin, SHS alone can be successful in characterizing membrane protein interactions, and the combination with single-channel current recordings provides a promising method for bio-nanotechnology. SHS - a direct, label-free, non-invasive technique - in conjunction with single-channel current recording can be used to screen pore variants with enhanced membrane incorporation sensitivity.

## MATERIALS AND METHODS

### A. Chemicals

Lipids 1,2-dioleoyl-*sn*-glycero-3-phosphocholine (DOPC), and 1,2-dioleoyl-*sn*-glycero-3-phospho-L-serine (sodium salt) (DOPS), were purchased in powder form (>99%) from Avanti Polar Lipids (Alabama, USA) and stored at −20 °C until further use. Chloroform for spectroscopy Uvasol^®^ (≥99%, Merck), methanol (≥99.9%, Fisher Chemical), sodium acetate (≥99%, Sigma-Aldrich), acetic acid (≥99.7%, Sigma-Aldrich), sodium phosphate dibasic (≥99%, Fluka), sodium phosphate monobasic (≥99%, Fluka), Trizma hydrochloride (≥99%, Sigma-Aldrich), Trizma base (≥99.9%, Sigma-Aldrich), and sodium chloride (≥99.999%, Acros) were used as received. Deconex 11 UNIVERSAL (Borer Chemie) was used as a cleaning solution. Water was purified by a Milli-Q UF-Plus instrument from Millipore, Inc., and it has an electrical resistivity of 18.2 MΩ⋅cm. All glassware was washed with a 5% deconex cleaning detergent solution in the ultrasonic bath for 30 min, then they were cleaned with Milli-Q ultrapure water in the sonication bath for another 20 min. After the cleaning, the glassware was rinsed with ultrapure water.

### B. Proaerolysin expression and purification

Proaerolysin wt and mutants were expressed using a pET22b vector, which allows periplasmic expression of the toxin and with a hexa-His tag on the C-terminus, as previously described^5,10^. Mutagenesis was carried out by using the QuickChange II XL kit (Agilent Technologies). Briefly, BL21(DE3)pLysS *E. coli* containing the wt or the mutant aerolysin expression plasmid were grown at 37°C up to an OD_600_ of 0.6-0.7. Isopropyl β-D-1-thiogalactopyranoside (IPTG) at a final concentration of 0.25 mM was added, and the temperature was decreased to 20°C for protein production overnight. Cells were harvested, when the OD_600_ reached 1.2, and resuspended in 20 mM sodium phosphate, 500 mM NaCl, pH 7.4 supplemented with cOmplete Protease Inhibitor cocktail (Roche). The cells were lysed by sonication and the supernatant was centrifuged at 12,000*rpm* for 35 min at 4°C, and loaded onto a HisTrap chelating column (GE Healthcare) running on an AKTA™ FPLC workstation. The protein was eluted in a 20 mM sodium phosphate buffer pH 7.4, 0.5 M NaCl buffer with a linear gradient of imidazole (0–0.5 M). Finally, fractions containing the protein were buffer-exchanged in 20 mM Tris, 500 mM NaCl, pH 7.4 by using a HiPrep Desalting column (GE Healthcare) before snap freezing and storing at −80°C.

### C. Sample preparation

Large unilamellar vesicles (LUVs) were prepared by the lipid film hydration method followed by extrusion. Lipid solutions were created by dissolving 10 mg of lipid powder in chloroform in a round-bottom glass tube. To evaporate the chloroform a gentle stream of N_2_ was directed into the rotating glass tube. The residual chloroform was dried under a room temperature vacuum for at least 3 h. The lipid film that was deposited on the glass wall was hydrated in 1 mL of ultrapure water that was heated up to above the respective phase-transition temperature of the used lipids. The resulting multilamellar vesicle solutions were extruded through a 100 nm diameter polycarbonate membrane in a Mini extruder (Avanti Polar Lipids). The LUVs were prepared in 50 *µ*M NaCl solution. LUVs were stored in closed containers for up to a week at 4 °C. The size of the resulting large unilamellar vesicles (LUVs) was determined by dynamic light scattering (DLS) using ZetaSizer Nano ZS (Malvern Instruments Ltd., UK). The z-average diameter of the vesicle is a result of three averaged measurements. The diameters of the DOPC LUVs are 118 nm with a PDI of 0.07 and their ζ-potential values are -2 ± 8 mV. The DOPC doped with 1% DOPS LUVs are 124 nm in diameter with a PDI of 0.07 and their ζ-potential values are -36 ± 16 mV. The concentration of total lipids was 0.5 mg/mL for SHS, DLS, and electrokinetic measurements. The ionic strength of the buffers and aerolysin stock solutions was kept the same for all the SHS, DLS, and electrophoretic measurements. The aqueous solution consisted of 50 µM NaCl, 13.3 µM MES buffer and 100 µM of buffer for pH adjustment. For pH between 4 and 5 Na acetate buffer was used, in the range from pH 6 to 7 phosphate buffer was used, and for the pH from 7.4 and 9 we used Trizma buffer. The pH of the solution was measured using a pH meter (HI 5522 pH/ISE/EC bench meter and HI 1330 pH electrode). The measured pH was always less than 0.16 points from the desired value. The sample volume used for SHS was 800 μL. Purified wt and mutants of aerolysin were activated as previously described^8^. Briefly, the toxin was diluted to the concentration of 0.2 μg/ml and then incubated at 4°C with Trypsin-agarose (Sigma-Aldrich Chemie GmbH, Buchs, SG Switzerland) for 2 hours to activate the toxin for oligomerization. The solution was centrifuged (10,000*g*, 4°C, 10min) to remove the trypsin-agarose beads. Activated toxins were aliquoted and kept at -80°C. Protein concentration was checked by Nanodrop and added accordingly.

### D. Second-harmonic scattering

Figure 1A shows the AR-SHS setup, already described in Ref 17. AR-SHS measurements were performed using 190-fs laser pulses centered at 1028 nm with a 200 kHz repetition rate. The polarization of input pulses was controlled by a Glan-Taylor polarizer (GT10-B, Thorlabs) in combination with a zero-order half-wave plate (WPH05M-1030). The filtered (FEL0750, Thorlabs) input pulses with a pulse energy of 0.3 μJ (incident laser power P = 60 mW) were focused into a cylindrical glass sample cell (inner diameter 4.2 mm) with a beam waist of 2ω_0_ ≈ 36 μm and a corresponding Rayleigh length of ∼0.94 mm. The scattered SH beam generated was analyzed (GT10-A, Thorlabs), filtered with a notch filter (ZET514/10x,Chroma), collimated with a plano-convex lens (f = 5 cm), and finally focused into a gated photomultiplier tube (H7421-40; Hamamatsu). The data points were acquired as an average of 100 measurements with a 1.5 s integration time and a gate width of 10 ns. The detection angle θ, which has an acceptance angle of 11.4°, was set to 45°, which corresponds to the angle of the maximum SH intensity. All measurements were performed in a temperature- and humidity-controlled room (T = 297 K; relative humidity, 26.0 %).

In a non-resonant SHS experiment, a pulsed femtosecond near-infrared laser beam interacts with the liposome solution. SH photons are emitted from all non-centrosymmetric molecules that are non-centrosymmetrically distributed. Since interfacial water is non-centrosymmetrically distributed while the bulk liquid is not, SHS has an exquisite interfacial sensitivity. When LUVs are dispersed in the water the orientation of water molecules will be perturbed at the interface of these particles. The interfacial water outnumbers lipids with a ratio of >1:100, and due to the non-resonant nature of the process, both constituents respond with electromagnetic fields that have equal magnitudes. SH intensity depends quadratically on the emitted electric field (i.e. number density of the molecules). Thus, the SHS signal generally reports on membrane hydration arising from the net orientational order of water molecules along the surface normal.

The normalized SH intensity at the angle was calculated as

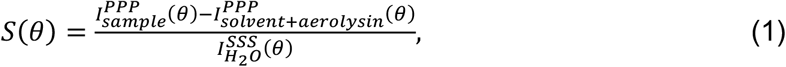

where 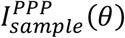 and 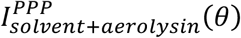 are the average SHS intensities of the sample (solvent, liposomes, and aerolysin) and solvent with aerolysin, respectively. PPP and SSS represent the polarization state of the outgoing and incident light relative to the scattering plane (P - parallel or S - perpendicular).

The normalized coherent SH intensity difference *Δ S* was calculated for each concentration of aerolysin as a difference between normalized SH intensity of LUVs without added aerolysin *S*_*L*_ (*θ*) and LUVs with the given concentration of aerolysin, *S*_*LP*._(*θ*), *Δ S* = |*S*_*L*_−*S*_*LP*._|. This value was then normalized to 1 for better comparison between the samples. The fitting procedure was previously described in more detail in Ref.^23^. The measured difference in the coherent SH intensity *Δ S* is

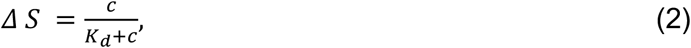

where *c* is the aerolysin concentration and *K*_*d*_ is the dissociation constant. The error bars were determined as a standard deviation from 100 measurements. As a result of the aerolysin concentration increase (adding volume), the particles in the solution were getting diluted so the linear correction for the number of particles in the focal spot was applied. In the dissociation constant calculations, we assumed each protein insertion was a unique event.

### E. Single-channel current recording experiments

Phospholipid of 1,2-dioleoyl-*sn*-glycero-3-phosphocholine powder (Avanti Polar Lipids Inc., Alabaster, AL, USA) was dissolved in octane (Sigma-Aldrich Chemie GmbH, Buchs, Switzerland) to a final concentration of 8 mg/mL. Purified wt and mutants of aerolysin were activated as previously described^8^. Briefly, the toxin was diluted to the concentration of 0.2 μg/ml and then incubated at 4°C with Trypsin-agarose (Sigma-Aldrich Chemie GmbH, Buchs, SG Switzerland) for 2 hours to activate the toxin for oligomerization. The solution was centrifuged (10,000*g*, 4°C, 10min) to remove the trypsin-agarose beads. Activated toxins were aliquoted and kept at -80°C. Nanopore single-channel current recording experiments were performed on an Orbit Mini instrument equipped with a temperature control setup (Nanion, Munich, Germany). Phospholipid membranes were formed across a MECA 4 recording chip that contains 4 circular microcavities (size 50 µm diameter) in a highly inert polymer. Each cavity contains an individual integrated Ag/AgCl microelectrode and can record four artificial lipid bilayers in parallel. The buffers for the different experiments include 10 mM phosphate buffer pH 7.4, 10 mM Tris pH 9, and 10 mM sodium acetate pH 5 all complemented with 1M KCl. The temperature was set to 25°C for all experiments. Once a stable lipid baseline with a capacitance also above 6 pF, the activated toxins were added and insertion(s) of the pore was recorded for each aerolysin mutants and recorded with Elements Data Reader (Elements srl, Italy) and further analyzed by using Clampfit (Axon, Molecular device). Experiments were repeated at least 3 times to obtain reproducibility. Results and graphs were produced in Origin (OriginLab Corporation) and figures and tables were generated in Adobe Illustrator 2022 (Adobe).

### F. Visualization and calculation of the protonation state of aerolysin

To calculate the protonation states of the aerolysin protein, we used the APBS web server^26^, where we prepared aerolysin in the heptamer oligomeric state (with PDB ID: 5JZT^27^) at different pH (by using PROPKA to assign the different protonation states at pH 4, 5, 6, 7, 8 and 9) by using the AMBER force field. Once the calculation of the protein was performed, the pqr files were loaded into PyMOL^28^ and the APBS Electrostatics^26^ plugin (in mg-auto configuration) was used to generate the electrostatic map for the aerolysin at different pH. The pictures were generated in PyMOL^28^, and the figure was created in Adobe Illustrator 2022 (Adobe).

## REFERENCES

(1) Dal Peraro, M.; van--der--Goot F.G. Pore-Forming Toxins: Ancient, but Never Really out of Fashion. Nat. Rev. Microbiol. 2016, 14 (2), 77–92. 10.1038/nrmicro.2015.3.

(2) Ulhuq, F. R.; Mariano, G. 2022. Bacterial Pore-Forming Toxins. Microbiology 168 (3), 001154. 10.1099/mic.0.001154.

(3) Rossjohn, J.; Feil S.C.; McKinstry, W. J.; Tsernoglou, D.; Van Der Goot, G.; Buckley J.T.; Parker, M.W. Aerolysin—A Paradigm for Membrane Insertion of Beta-Sheet Protein Toxins? J. Struct. Biol. 1998, 121 (2), 92–100. 10.1006/jsbi.1997.3947.

(4) Cirauqui, N.; Abriata, L. A.; van--der--Goot, F. G.; Dal--Peraro, M. Structural, Physicochemical and Dynamic Features Conserved within the Aerolysin Pore-Forming Toxin Family. Sci. Rep. 2017, 7 (1), 13932. 10.1038/s41598-017-13714-4.

(5) Iacovache, I.; Degiacomi M.T.; Pernot, L.; Ho, S.; Schiltz, M.; Peraro, M. D.; van--der--Goot, F.G. Dual Chaperone Role of the C-Terminal Propeptide in Folding and Oligomerization of the Pore-Forming Toxin Aerolysin. PLOS Pathog. 2011, 7 (7), e1002135. 10.1371/journal.ppat.1002135.

(6) Degiacomi, M. T.; Iacovache, I.; Pernot, L.; Chami, M.; Kudryashev, M.; Stahlberg, H.; van--der--Goot, F. G.; Dal Peraro, M. Molecular Assembly of the Aerolysin Pore Reveals a Swirling Membrane-Insertion Mechanism. Nat. Chem. Biol. 2013, 9 (10), 623–629. 10.1038/nchembio.1312.

(7) Tsitrin, Y.; Morton C.J.; El Bez, C.; Paumard, P.; Velluz, M.-C.; Adrian, M.; Dubochet, J.; Parker, M. W.; Lanzavecchia, S.; van der Goot, F. G. Conversion of a Transmembrane to a Water-Soluble Protein Complex by a Single Point Mutation. Nat. Struct. Biol. 2002, 9 (10), 729–733. 10.1038/nsb839.

(8) Cao, C.; Cirauqui, N.; Marcaida, M. J.; Buglakova, E.; Duperrex, A.; Radenovic, A.; Dal Peraro, M. Single-Molecule Sensing of Peptides and Nucleic Acids by Engineered Aerolysin Nanopores. Nat. Commun. 2019, 10 (1), 4918. 10.1038/s41467-019-12690-9.

(9) Cressiot, B.; Ouldali, H.; Pastoriza-Gallego, M.; Bacri, L.; Van der Goot, F. G.; Pelta, J. Aerolysin, a Powerful Protein Sensor for Fundamental Studies and Development of Upcoming Applications. ACS Sens. 2019, 4 (3), 530–548. 10.1021/acssensors.8b01636.

(10) Cao, C.; Li, M.-Y.; Cirauqui, N.; Wang, Y.-Q.; Dal Peraro, M.; Tian, H.; Long, Y.-T. Mapping the Sensing Spots of Aerolysin for Single Oligonucleotides Analysis. Nat. Commun. 2018, 9 (1), 2823. 10.1038/s41467-018-05108-5.

(11) Fennouri, A.; Przybylski, C.; Pastoriza-Gallego, M.; Bacri, L.; Auvray, L.; Daniel, R. Single Molecule Detection of Glycosaminoglycan Hyaluronic Acid Oligosaccharides and Depolymerization Enzyme Activity Using a Protein Nanopore. ACS Nano 2012, 6 (11), 9672–9678. 10.1021/nn3031047.

(12) Piguet, F.; Ouldali H.; Pastoriza-Gallego, M.; Manivet, P.; Pelta, J.; Oukhaled, A. Identification of Single Amino Acid Differences in Uniformly Charged Homopolymeric Peptides with Aerolysin Nanopore. Nat. Commun. 2018, 9 (1), 966. 10.1038/s41467-018-03418-2.

(13) Ouldali, H.; Sarthak K.; Ensslen, T.; Piguet, F.; Manivet, P.; Pelta, J.; Behrends, J. C.; Aksimentiev, A.; Oukhaled, A. Electrical Recognition of the Twenty Proteinogenic Amino Acids Using an Aerolysin Nanopore. Nat. Biotechnol. 2020, 38 (2), 176–181. 10.1038/s41587-019-0345-2.

(14) Afshar BakshlooM; Kasianowicz, J. J.; Pastoriza-Gallego, M.; Mathé, J.; Daniel, R.; Piguet, F.; Oukhaled, A. Nanopore-Based Protein Identification. J. Am. Chem. Soc. 2022, 144 (6), 2716–2725. 10.1021/jacs.1c11758.

(15) Bhatti, H.; Jawed R.; Ali, I.; Iqbal, K.; Han, Y.; Lu, Z.; Liu, Q. Recent Advances in Biological Nanopores for Nanopore Sequencing, Sensing and Comparison of Functional Variations in MspA Mutants. RSC Adv. 11 (46), 28996–29014. 10.1039/d1ra02364k.

(16) Lu, S.-M.; Wu X.-Y.; Li, M.-Y.; Ying, Y.-L.; Long, Y.-T. Diversified Exploitation of Aerolysin Nanopore in Single-Molecule Sensing and Protein Sequencing. VIEW 2020, 1 (4), 20200006. 10.1002/VIW.20200006.

(17) Gomopoulos, N.; Lütgebaucks, C.; Sun, Q.; Macias-Romero, C.; Roke, S. Label-Free Second Harmonic and Hyper Rayleigh Scattering with High Efficiency. Opt. Express 2013, 21 (1), 815. 10.1364/OE.21.000815.

(18) Lütgebaucks, C.; Gonella, G.; Roke, S. Optical Label-Free and Model-Free Probe of the Surface Potential of Nanoscale and Microscopic Objects in Aqueous Solution. Phys. Rev. B 2016, 94 (19). 10.1103/PhysRevB.94.195410.

(19) Gonella, G.; Lütgebaucks, C. Second Harmonic and Sum Frequency Generation from Aqueous Interfaces Is Modulated by Interference. 2.

(20) Schönfeldová, T.; Piller, P.; Kovacik, F.; Pabst, G.; Okur, H. I.; Roke S. Lipid Melting Transitions Involve Structural Redistribution of Interfacial Water. J. Phys. Chem. B 2021, 125 (45), 12457–12465. 10.1021/acs.jpcb.1c06868.

(21) Dedic, J.; Rocha, S.; Okur, H. I.; Wittung-Stafshede, P.; Roke, S. Membrane–Protein– Hydration Interaction of α-Synuclein with Anionic Vesicles Probed via Angle-Resolved Second-Harmonic Scattering. J. Phys. Chem. B 2019, 123 (5), 1044–1049. 10.1021/acs.jpcb.8b11096.

(22) Roesel, D.; Eremchev, M.; Schönfeldová, T.; Lee, S.; Roke, S. Water as a Contrast Agent to Quantify Surface Chemistry and Physics Using Second Harmonic Scattering and Imaging: A Perspective. Appl. Phys. Lett. 2022, 120 (16), 160501. 10.1063/5.0085807.

(23) Schönfeldová, T.; Okur, H. I.; Vezočnik, V.; Iacovache, I.; Cao, C.; Dal Peraro, M.; Maček, P.; Zuber, B.; Roke, S. Ultrasensitive Label-Free Detection of Protein– Membrane Interaction Exemplified by Toxin-Liposome Insertion. J. Phys. Chem. Lett. 2022, 13 (14), 3197–3201. 10.1021/acs.jpclett.1c04011.

(24) Iacovache, I.; Paumard P.; Scheib, H.; Lesieur, C.; Sakai, N.; Matile, S.; Parker, M. W.; van der Goot, F. G. A Rivet Model for Channel Formation by Aerolysin-like Pore-Forming Toxins. EMBO J. 2006, 25 (3), 457–466. 10.1038/sj.emboj.7600959.

(25) Sauciuc, A., Morozzo della Rocca, B., Tadema, M.J.., et al. Translocation of linearized full-length proteins through an engineered nanopore under opposing electrophoretic force. Nat. Biotechnol. 2023, 10.1038/s41587-023-01954-x

(26) Jurrus, E.; Engel, D.; Star, K.; Monson, K.; Brandi, J.; Felberg, L. E.; Brookes, D. H.; Wilson, L.; Chen, J.; Liles, K.; Chun, M.; Li, P.; Gohara, D. W.; Dolinsky, T.; Konecny, R.; Koes, D. R.; Nielsen, J. E.; Head-Gordon, T.; Geng, W.; Krasny, R.; Wei, G.-W.; Holst, M. J.; McCammon, J. A.; Baker, N. A. Improvements to the APBS Biomolecular Solvation Software Suite. Protein Sci. 2018, 27 (1), 112–128. 10.1002/pro.3280.

(27) Iacovache, I.; De Carlo S.; Cirauqui, N.; Dal Peraro, M.; van der Goot, F. G.; Zuber, B. Cryo-EM Structure of Aerolysin Variants Reveals a Novel Protein Fold and the Pore-Formation Process. Nat. Commun. 2016, 7, 12062. 10.1038/ncomms12062.

(28) The PyMOL Molecular Graphics System.

