## Supporting information for "Dissecting the membrane association mechanism of aerolysin pore at femtomolar concentrations using water as a probe"

### Dissecting the membrane association mechanism of the aerolysin pore by high-throughput second harmonic scattering

#### SHS patterns for pH 4 and pH 9

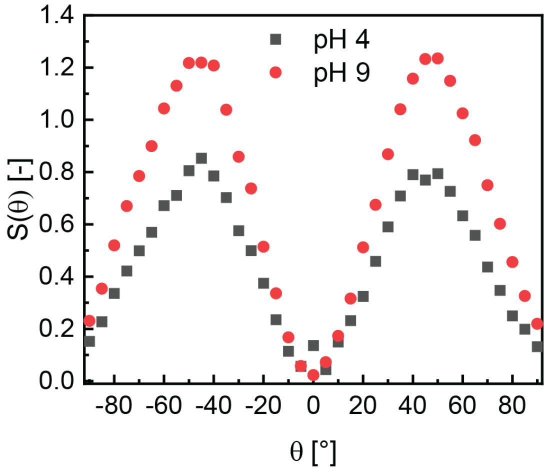


**Figure S1:** SHS pattern of LUVs consisting of DOPC with 1 mol % of DOPA with 5 x $10^{-10}$ M wt aerolysin at pH 4 and 9. The maximum SH intensity occurs around $\theta$ = 45 °. We used 1 mol % of DOPA to increase the signal to noise ratio for these measurements.
